# GRep: Gene Set Representation via Gaussian Embedding

**DOI:** 10.1101/519033

**Authors:** Sheng Wang, Emily Flynn, Russ B. Altman

## Abstract

Molecular interaction networks are our basis for understanding functional interdependencies among genes. Network embedding approaches analyze these complicated networks by representing genes as low-dimensional vectors based on the network topology. These low-dimensional vectors have recently become the building blocks for a larger number of systems biology applications. Despite the success of embedding genes in this way, it remains unclear how to effectively represent gene sets, such as protein complexes and signaling pathways. The direct adaptation of existing gene embedding approaches to gene sets cannot model the diverse functions of genes in a set. Here, we propose GRep, a novel gene set embedding approach, which represents each gene set as a multivariate Gaussian distribution rather than a single point in the low-dimensional space. The diversity of genes in a set, or the uncertainty of their contribution to a particular function, is modeled by the covariance matrix of the multivariate Gaussian distribution. By doing so, GRep produces a highly informative and compact gene set representation. Using our representation, we analyze two major pharmacogenomics studies and observe substantial improvement in drug target identification from expression-derived gene sets. Overall, the GRep framework provides a novel representation of gene sets that can be used as input features to off-the-shelf machine learning classifiers for gene set analysis.

## 1. INTRODUCTION

Molecular interaction networks provide novel insights into the functional interdependencies among genes and proteins[1,2]. In particular, recently developed high-throughput experimental techniques, such as yeast two-hybrid screens and genetic interaction assays, have enabled researchers to piece together large-scale interaction networks in bulk[3,4]. Consequently, a variety of network-based approaches, including network propagation[5–11], network clustering[12,13], network integration[14,15], and network regularization[16], have been developed to efficiently analyze these networks. Among them, network embedding has emerged as a powerful network analysis approach because it generates a highly informative and compact vector representation for each node in the network[15,17,18]. Molecular interaction networks are noisy and incomplete, especially as they increase in size[15,18]. Network embedding adapts dimensionality reduction techniques to de-noise and improve accuracy in high-dimensional network data. In addition, before the advent of network embedding approaches, researchers identified network features for machine learning by hand, which is time-consuming and often requires expert knowledge. By contrast, network embedding automates this process by representing each node in the network as a vector. These node representations have shown good performance in machine learning classifiers, and thus become building blocks of a large number of systems biology applications[15,18–21]. Throughout this paper, we use genes for nodes; however, it is important to note that these methods can be applied to any type of node.

Biologically meaningful gene sets provide useful prior knowledge about how genes may work together. There are a huge number of publicly available gene sets[22–25], which come from many sources: Genome-wide association studies (GWAS) produce sets of genes associated with a disease or other phenotype. Gene expression analyses identify gene sets by examining differential expression between conditions, or clustering genes by expression similarity. Biological network analysis implicates genes, proteins, or metabolites that interact with one another in the same network neighborhoods. In all of these cases, the hypothesis is that genes in the set are involved in the same biological processes or functions. Moreover, these gene sets have emerged as useful prior knowledge to boost signal-to-noise and increase explanatory power. Consequently, biologically meaningful gene sets have been involved in a large number of prediction tasks, such as drug-pathway interaction[26] and disease signature prediction[27]. However, a substantial bottleneck for gene set-based analysis is a lack of good feature representations for those gene sets. Because gene sets are widely used in bioinformatics analyses and provide important signal, learning highly informative and compact representations for gene sets has the potential to improve a large number of clinical and biological applications.

While many useful gene representations are now available[15,17,18], learning a representation for a gene set remains challenging. The arguably simplest approach to represent a gene set is the average of its individual gene representations. In natural language processing, naively averaging word vectors has been successfully used to construct sentences embedding[28]. However, in contrast with sentences which only have a few words, gene sets can be arbitrarily large. Average embedding s then not expressive enough to represent such a gene set. Fig. 1a shows an example where average embedding approach is not able to distinguish two completely different gene sets. Another simple approach to represent a gene set is to add new “gene set nodes” to the existing molecular network and connect them to their gene members. One can then run node embedding on this network to obtain the representation of each gene set. However, adding these potentially high-degree nodes to the network can substantially change the topological structure of the network, leading to an inaccurate estimation of the embedding space. Other methods aim at embedding densely connected subnetworks: ComE performs community embedding and community detection simultaneously using a community-aware high-order proximity[29]. PathEmb models pathways as documents and then applies document embedding models to calculate pathway similarity[30]. However, both ComE and PathEmb require gene sets to be connected in the network, which is not the case for most of the biologically meaningful gene sets.

**Figure 1.**
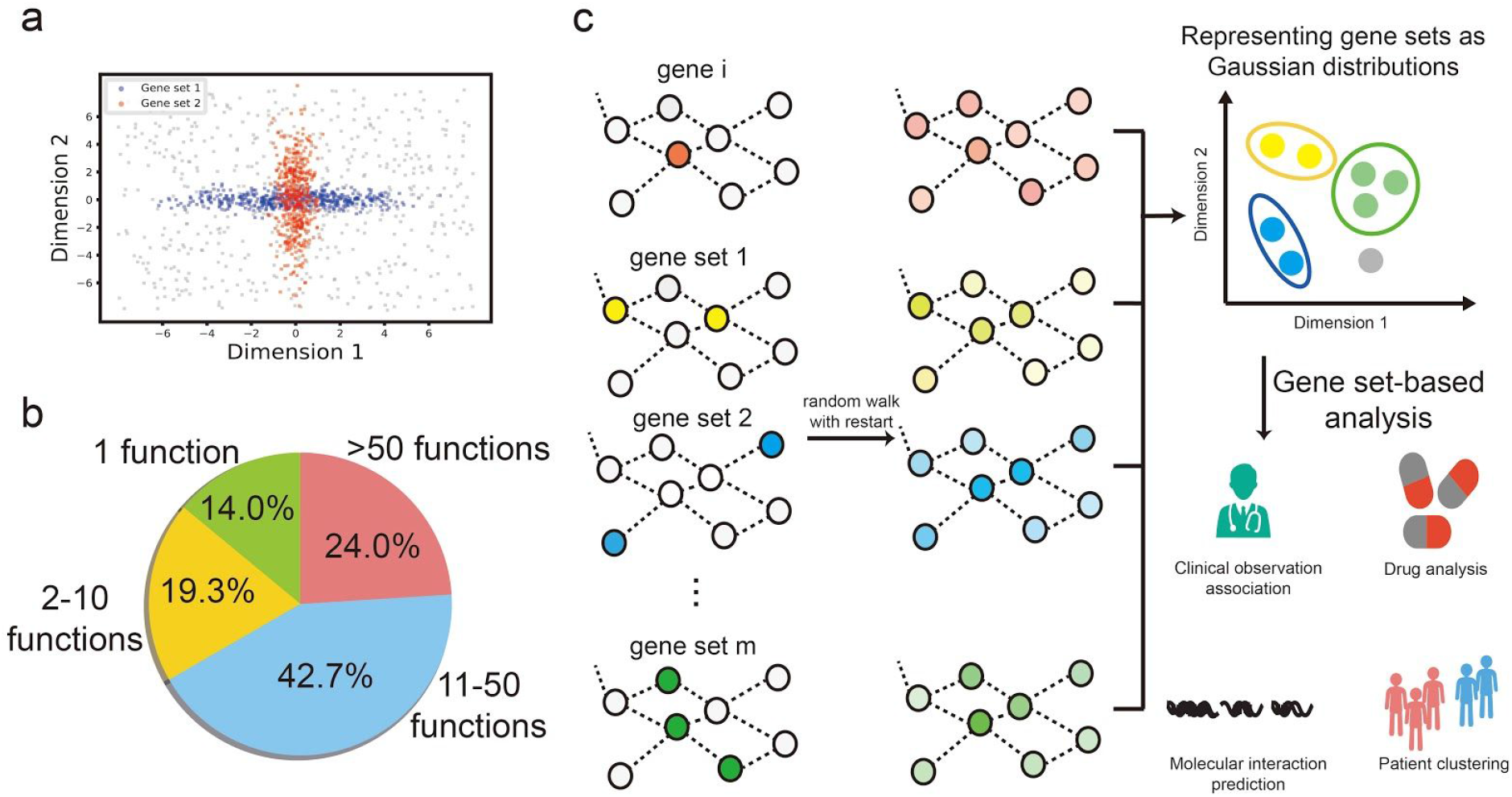
a. Two different gene sets are embedded in the same point (0,0) if we simply average the embeddings of individual genes. b. Gene sets contain a variety of functions; this pie chart shows the percent of significantly enriched Gene Ontology functions in 150 drug response correlated gene sets (P < 0.05 after Bonferroni correction). c. Flowchart describing GRep embedding process and downstream applications: RWR is used to calculate the diffusion states of each gene and gene set. These diffusion states are then embedded in a low-dimensional space where genes are represented as single points and gene sets are represented as Gaussian distributions. These representations can be applied to a variety of gene set-based analysis.

Moreover, all of these approaches rely on the assumption that genes in the same set tend to have similar properties. This is intuitively the case for protein complexes or biological processes; however, this is not true for a large number of other biologically meaningful gene sets. To examine the diversity of functions in gene sets, we calculated the Gene Ontology enrichment of 150 drug response-related gene sets derived from two pharmacogenomics datasets (Fig. 1b)[31]. We found that 86% of these gene sets are significantly enriched with more than one function (P<0.05 after Bonferroni correction). Genes in the same set may still have different functions or be involved in different biological processes; however, this diversity is ignored in simple average embedding. Recently, Gaussian embedding, which represents each node as a multivariate Gaussian distribution in the low-dimensional space, has been extensively used to model the uncertainty of nodes [32–34]. Despite its success in representing genes, Gaussian embedding has not yet been applied to gene sets, which are more diverse than individual genes. Motivated by prior work on Gaussian embedding, we propose to represent each gene set as a multivariate Gaussian distribution according to its network topology. When using an embedding vector to represent a gene set, the diversity can be modeled by the uncertainty of each dimension in this vector. Dimensions that have small variance across different genes in the set should be more reliable. Therefore, a more robust approach to represent a gene set is to use two parameters: one represents the average values in each dimension, and the other represents the uncertainty of each dimension. These two parameters define the mean and the covariance matrix of a multivariate Gaussian distribution. To the best of our knowledge, our method is the first approach that learns compact representations for gene sets.

In this paper, we present GRep (Gene set Representation), a novel computational method that represents each gene set as a highly informative and compact multivariate Gaussian distribution (Fig. 1c). GRep takes a biological network and a collection of gene sets as input. It represents each gene as a single point and each gene set as a multivariate Gaussian distribution parameterized by a low-dimensional mean vector and a low-dimensional covariance matrix. The mean vector of each gene set describes the joint contribution of genes in this gene set, and the covariance matrix characterizes the agreement among individual genes in each dimension. By using this representation, GRep is able to differentiate between gene sets that would be considered equivalent by average embedding. The key idea of GRep is to use the prior knowledge in gene sets and group genes in the same set closely as a multivariate Gaussian distribution in the low-dimensional space. To achieve this, GRep solves an optimization problem to preserve the network topology according to diffusion states. We evaluate GRep on a collection of drug response correlated gene sets derived from Genomics of Drug Sensitivity in Cancer (GDSC)[35] and The Cancer Therapeutic Portal (CTRP)[36]. We demonstrate that representing those gene sets using GRep substantially outperforms comparison approaches on drug-target identification in both datasets.

## 2. METHODS

Biologically meaningful gene sets, such as signaling pathways and protein complexes, aggregate gene level information into higher level patterns. A key observation behind our approach is that gene sets can have diverse molecular functions and/or biological processes. GRep explicitly models this diversity as a low-dimensional Gaussian distribution which summarizes both location and uncertainty of each dimension. To summarize, GRep takes a network and a collection of gene sets as input (Fig. 1c). It first calculates the diffusion states of each gene and gene set to characterize their topological information in the network. GRep then finds the low-dimensional representations for genes and gene sets according to these diffusion states. Each gene is represented as a single point in the low-dimensional space. Each gene set is represented as a multivariate Gaussian distribution which is parameterized by a mean vector and a covariance matrix.

**Problem definition.** Let *A* ∈ *R^n×n^* be the adjacency matrix of a given network *G*, where *n* is the number of genes. *V* denotes the set of all genes. Let *H* = {*h*_1_,*h*_2_,…,*h_m_*} be *m* gene sets defined on *G*, where each gene set *h_i_* is a set of genes 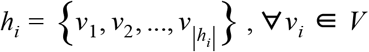. GRep aims to find a low-dimensional multivariate Gaussian distribution *N*(*μ_h_*,Σ_*h*_) for each gene set *h* with mean *μ_h_* ∈ *R^d^* and covariance matrix *Σ_h_* ∈ *R^d×d^*, where *d*≪*n*.

### Random walk with restart from a gene set

In order to define the objective function, we first need to characterize the network topology that we want to preserve in the low-dimensional space. Here, we use random walk with restart (RWR) to capture the network topology. RWR captures fine-grain topological properties that lie beyond direct neighbors[5,6]. When there are missing and spurious genes in a given gene set, RWR can correct the noise using network neighbors. RWR differs from the conventional random walk in that it introduces a predefined probability of restarting at the initial gene after every iteration.

Formally, we first calculate a transition matrix *B*, which represents the probability of a transition from gene *i* to gene *j*. *B* is defined as:

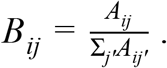

To run RWR from gene *i*, we define 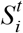 as an *n*-dimensional distribution vector in which each entry *j* contains the probability of gene *j* being visited from gene *i* after *t* steps. RWR from gene *i* with restart probability *p_S_* is defined as:

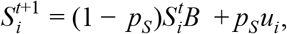

where *u_i_* is an *n*-dimensional distribution vector with *u_i_*(*i*) = 1 and *u_i_*(*j*) = 0, *∀j* ≠ *i*. We can obtain the stationary distribution 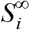 of RWR at the fixed point of this iteration. Consistent with the previous work[5,7,15,18], we define the diffusion state 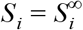 of each gene *i* to be the stationary distribution of an RWR starting at each gene. Here, the restart probability controls the relative influence of global and local topological information in the diffusion, where a larger value places greater emphasis on the local structure.

To run RWR from gene set *k*, we define 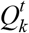 as an *n*-dimensional distribution vector in which each entry contains the probability of a gene being visited from gene set *k* after *t* steps. RWR from gene set *k* with restart probability *P_Q_* is defined as:

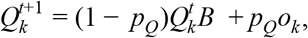

where *o_k_* is an *n*-dimensional distribution vector with 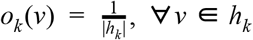 and 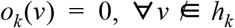. We can obtain the stationary distribution 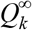 of RWR at the fixed point of this iteration. we define the diffusion state 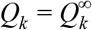 of each gene set *k* to be the stationary distribution of an RWR starting at each gene in *k* uniformly. When genes in the set are rank-ordered by importance, we can adjust *o_k_* according to the gene weights.

Notably, a gene set could have missing or spurious genes. RWR can account for the noisy gene sets using network neighbors to characterize the network topology. The restart probability reflects our uncertainty of this gene set, where a smaller value encourages the gene set to extend its members with network neighbors. GRep uses the diffusion state *S_i_* (*Q_k_*) to represent the topological information of gene *i* (gene set *k*) in the network. The *j* th entry *S_ij_* (*Q_kj_*) stores the probability that RWR starts at gene *i* (gene set *k*) and ends up at gene *j* in equilibrium.

### Representing gene sets as multivariate Gaussian distributions

The diffusion states of each gene and each gene set are then used to find the low-dimensional representation. GRep embeds genes and gene sets in the same low-dimensional space, where each gene is represented as a single point and each gene set is represented as a multivariate Gaussian distribution parameterized by a mean vector and a covariance matrix.

GRep uses two criteria to find the low-dimensional representation: 1) genes with similar diffusion states should be close to each other in the low-dimensional space, and 2) genes in a given gene set in the network should have higher probabilities in the Gaussian distribution of that gene set. The first criterion preserves the distance between genes and has been widely used in conventional node embedding approaches. The second criterion is unique to GRep, and groups genes in the same set together as a multivariate Gaussian distribution. By using the second criteria, GRep explicitly leverages the fact that genes in the same set are likely to have similar properties and thus should be closely located in the low-dimensional space.

Formally, let *L_gene_* and *L_set_* represent the loss function based on the above two criteria. The loss function can be defined as:

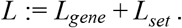

To preserve the gene distance (criteria 1), we define *L_gene_* as:

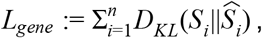

where *D_KL_* is the Kullback-Leibler (KL) divergence and 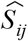 is defined as:

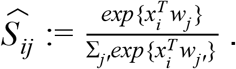

Here, *x_i_* is the representation of gene *i* in the low-dimensional space and *w_i_* is the context feature describing the network topology of gene *j*. If *x_i_* and *w_j_* are close in direction and have a large inner product, then it is likely that *j* is frequently visited in the random walk restarting from gene *i*. We optimize over *w* and *x* for all genes, using KL divergence as the objective function.

Similar to previous work, we relax the constraint that the entries in 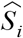 sum to one by dropping the normalization factor in the above equation[15,18]. As a result, 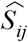 can be simplified as:

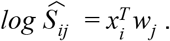

This simplification substantially reduces the computational complexity while still achieving comparable performance[15,18]. Since 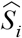 is no longer an *n*-dimensional probability simplex, we use the sum of squared errors instead of KL divergence as the new objective function.

Therefore, *L_gene_* is defined as:

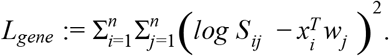

Next, to preserve the distance between genes and gene sets, we define *L_set_* as:

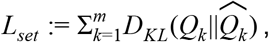

where 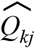 is defined as:

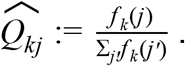

*f_k_* is the multivariate Gaussian probability density function and *f_k_*(*j*) is the probability density of gene *j*:

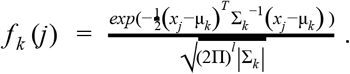

Here, we can optimize over the mean vector μ_*k*_ and the covariance matrix Σ_*k*_ to obtain the multivariate Gaussian distribution of gene set *k*.

Same as the simplification in *L_gene_*, we also drop the normalization factor in the above equation. As a result, 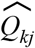 is simplified as:

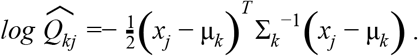

Notably, 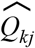 can also be viewed as the Mahalanobis distance of gene *j* from the mean μ_*k*_ and covariance matrix Σ_*k*_. The Mahalanobis distance can account for different variances in each direction and reduces to Euclidean distance when Σ is an identity matrix. While matrix factorization approaches, such as singular value decomposition (SVD), also calculate a diagonal matrix Σ, GRep improves on this by optimizing different Σ_*k*_ for each gene set *k* in order to model the uncertainty of each gene set differently.

We then use the sum of squared errors as the objective function:

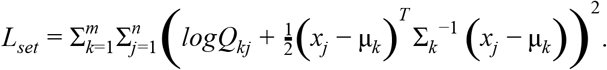

Summing up two parts, the new loss function of our model is defined:

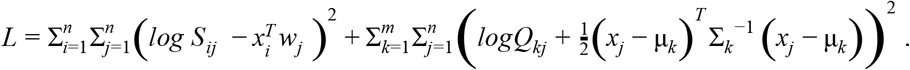

While the first term preserves gene distance in the network, the second term forces genes in the same set to form a multivariate Gaussian distribution. Therefore, these biologically meaningful gene sets are used as prior knowledge by GRep to infer the embedding of genes. By contrast, other methods, such as average embedding, are unable to leverage this prior knowledge.

### Parameter estimation

GRep has the following parameters: μ, Σ, *x*, and *w*. The parameters μ, *x*, and *w* can be directly estimated with gradient descent. By contrast, since Σ is the covariance matrix of a multivariate Gaussian distribution, we need to assure that it is positive semi-definite. To achieve this, let Λ_*k*_ be the precision matrix of the multivariate Gaussian distribution for gene set *k*:

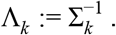

Instead of directly estimating the covariance matrix Σ_*k*_, we estimate the precision matrix Λ_*k*_ to avoid numerical problems that arise in matrix inversion. We define *C_k_* ∈ *R^d×d^* to force Λ_*k*_ to be positive semi-definite:

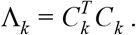

Since a matrix multiplied by its transpose is positive semi-definite, Λ_*k*_ is thus a positive semi-definite matrix. This further ensures that its inverse Σ_*k*_ is also a positive semi-definite matrix. Since there is no constraint on *C_k_*, we can use gradient descent to estimate *C_k_*. However, directly optimizing over *C_k_* introduces a substantial memory complexity of *O*(*md*^2^), which counteracts a key benefit of using a low-dimensional representation. To address this problem, we propose to factorize *C_k_* by using a low-rank approximation as:

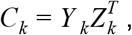

where 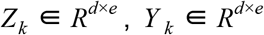, and *e*≪*d*. In our experiment, we found that set *e* to 5 is enough to obtain a good performance. The parameters of GRep are now μ, *Z*, *Y*, *x*, and *w*. We estimate these parameters using Adam to find a local optimum of this optimization problem[37].

After finding the low-dimensional representation of genes and gene sets in GRep, we can calculate the distance between gene sets and genes in the low-dimensional space. The distance 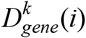 between gene *i* and gene set *k* is calculated according to the probabilistic density function of the multivariate Gaussian distribution for gene set *k*:

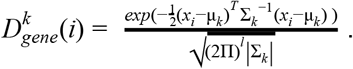

Using this formulation, the distance between a gene and a gene set depends not only on the mean vector μ (the location of this gene set) but also on the covariance matrix Σ. To calculate the distance 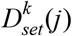 between gene set *k* and gene set *j*, we take the average asymmetric KL divergence according to their Gaussian distributions:

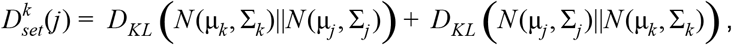

where 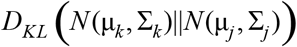 is calculated as:

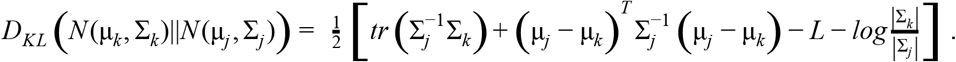

## 3. RESULTS

### Network and gene set collection

We evaluated GRep using the molecular interaction network from InBioMap and a collection of drug response correlated gene sets from expression data. InBioMap is a publicly available protein-protein interaction (PPI) network that aggregates PPIs from eight different gene orthology databases. Human protein pairs only are connected if the majority of the databases agree on the phylogenetic relationship between two proteins in model organisms or humans. The InBioMap network contains 15,108 genes and 3,621,168 edges. All edges are used as unweighted and undirected in our model.

To obtain drug response correlated gene sets, we need to collect both drug response data and gene expression data. We obtained two large-scale drug response screens from Genomics of Drug Sensitivity in Cancer (GDSC)[35] and The Cancer Therapeutics Response Portal (CTRP)[36]. Those two datasets are two of the existing largest pharmacogenomics studies and have been widely used to evaluate various pharmacogenomics analyses. We collected the gene expression of cell lines in these two studies from GDSC and Cancer Cell Line Encyclopedia (CCLE)[38], respectively. For each drug, we formed a set of genes whose expression is most correlated with response. We referred to this set of genes as response correlated gene set (RCGS). We used the absolute Spearman correlation coefficient larger than 0.4 as the criteria to form the gene set for each drug. We obtained 55 gene sets for GDSC and 175 gene sets for CTRP. Finally, we obtained the target of each drug from GDSC and CTRP.

### Experimental setting

To compare GRep with other approaches, we ask if the low-dimensional representations of RCGS can accurately identify drug targets. We hypothesize that drug targets will be close to response correlated genes in the network. Hence, a good gene set representation approach will place a RCGS of a given drug closely to its drug target in the low-dimensional space. We calculate the distance between the RCGS and all test genes to get a ranked list of genes for each drug. Then we measure the extent to which GSDC and CTRP true drug-target associations are concentrated near the top of the list using the area under the receiver operating characteristic curve (AUROC)[20].

Since there are no existing gene set embedding approaches for comparison, we propose six competitive comparison approaches adapted from the state-of-the-art node embedding approach as representative baselines. 1) Plain average embedding (**Plain avg**): each gene set is represented by the average of its gene embedding vectors. 2) Weighted average embedding of genes in the set (**Weighted avg set**): gene sets are represented by the weighted average of gene embedding vectors. We use diffusion states as weights here. 3) Weighted average embedding of all genes (**Weighted avg all**): Each gene set is represented as the weighted average of the gene embedding vectors of all genes in the network, with diffusion states as weights. 4) Heterogeneous network embedding (**Het emb**): We first construct a new heterogeneous network of genes and gene sets. Gene sets are added as new “gene set nodes” into the original gene network. An edge is constructed between a “gene set node” and a gene if the gene is in this gene set. We then run the node embedding approach on this new heterogeneous network to find the representation of each gene set. 5) Heterogeneous network decomposition (**Het SVD**): We use singular value decomposition (SVD) to decompose the adjacency matrix of the heterogeneous network we constructed above. 6) Random walk with restart (**RWR**): We use the diffusion state to represent each gene and gene set. These baselines cover the most competitive gene set embedding approaches we can think of.

We use diffusion component analysis (DCA), a recently developed node embedding algorithm, as the underlying node embedding approach for these baselines[15,18]. We use cosine similarity to calculate the proximity of a gene set and a gene in the low-dimensional space as suggested by DCA[15,18]. For each baseline, we iterate over a range of hyperparameters and select the best performing result. For our method, we set *e* to 5, *d* to 100, *p_Q_* to 0.5 and *p_S_* to 0.5. We observe that the performance of our method is not sensitive to these hyperparameters.

### GRep substantially improves drug-target identification

To evaluate GRep, we performed large-scale drug target identification on two pharmacogenomics studies, GDSC and CTRP. The results are summarized in Figure 2. Our approach significantly outperforms plain avg, weighted avg all and weighted avg set on both datasets. In CTRP, our method achieved 0.8667 AUROC, which is much higher than 0.7102 AUROC of plain avg, 0.7104 of weighted set avg and 0.7319 AUROC of weighted avg all. We noted that weighted avg all performs consistently better than plain avg and weighted avg set at this task. This suggests that gene sets could be noisy, and using diffusion states to smooth this gene set with network neighbors can substantially reduce the noise. The same improvement was observed on GDSC where our method achieved 0.8890 AUROC, which is again substantially higher than 0.6870 AUROC of plain avg, 0.7325 of weighted avg all and 0.6870 AUROC of weighted avg set. All improvements were statistically significant (P<0.05; pairs Wilcoxon signed-rank test). The above results suggest that representing a gene set through simple averaging is not able to modeling uncertainty, leading to worse performance. By incorporating prior knowledge about gene sets and jointly optimizing the gene and gene set representations, our method substantially improved drug target identification.

**Figure 2.**
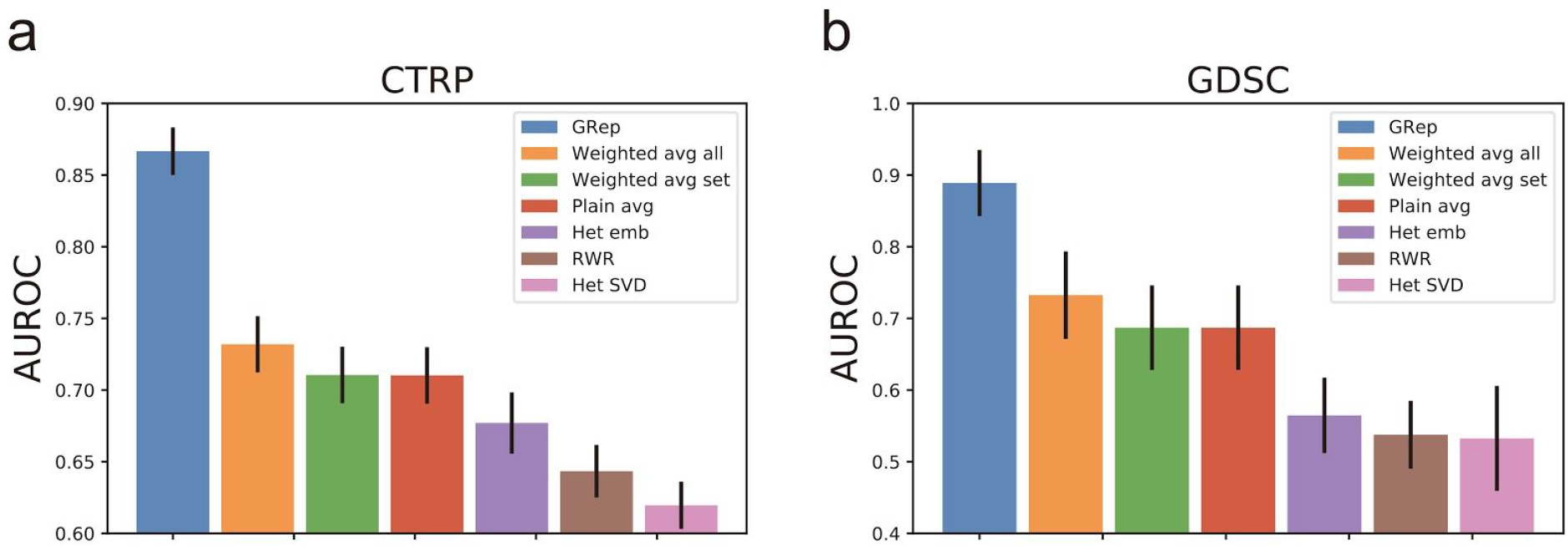
Performance of different gene set embedding methods on drug-target identification in CTRP (a) and GDSC (b).

To assess the effect of using a Gaussian distribution rather than single point representation, we compare GRep with three heterogeneous network-based approaches. All three approaches represent gene sets as single points, thus are unable to model the diverse function within each gene set. We found that that GRep substantially outperforms these three approaches on both datasets. For example, in CTRP, our method achieved 0.0.8667 AUROC, which is much higher than 0.6196 AUROC of Het SVD, 0.6434 AUROC of RWR and 0.6770 AUROC of Het emb. Similar to previous work [15], we observed that Het emb consistently outperforms Het SVD and RWR on this task. The poor performance of RWR could be due to the noisy diffusion state caused by missing or spurious edges in the network. All of the improvements were statistically significant (P<0.05; pairs Wilcoxon signed-rank test).

Interestingly, we found that heterogeneous network-based approaches are worse than averaging embedding on both datasets. Network embedding approaches rely on finding similar contexts (e.g., similar neighbors or similar diffusion states) to accurately embed different nodes. However, in a heterogeneous network, a “gene set node” could have a large number of neighbors and the neighborhood structure might be noisy, which may make it difficult to find enough other nodes with similar contexts to support an accurate embedding. Constructing a heterogeneous network may also introduce too many high degree nodes which substantially change the topological structure of the network.

## CONCLUSION

In this paper, we introduced GRep, a novel analytical method for learning gene set representation. To our knowledge, this is the first method for gene set embedding. GRep uses a multivariate Gaussian distribution to represent each gene set in order to model the diversity of genes in the same set. GRep leverages the prior knowledge that genes from the same set should have similar properties and thus be closely located in the low-dimensional space. In addition to localizing nodes in the low-dimensional space, GRep also captures the uncertainty of each dimension, which is not achieved by conventional approaches. Because there are no existing methods for gene set embedding, we constructed six competitive baselines by adapting conventional gene embedding approaches. GRep significantly outperforms these conventional approaches on a drug-target identification task in two large-scale pharmacogenomics studies.

In the future, we plan to pursue further improvements in drug-target identification with GRep, by incorporating other data such as somatic mutation and loss-of-function screens. We hypothesize that GRep will be substantially improved by training on a large collection of biologically meaningful gene sets simultaneously. In such a case, we want to use GRep to classify biologically meaningful gene sets and randomly generated gene sets. More importantly, while we focus on gene set analysis in this paper, the GRep framework is not limited to gene set analysis and can be applied to other biological set and biological network analysis, such as drug network and disease network.

The GRep software package is available at https://github.com/wangshenguiuc/GRep.

